# Dynamic synergetic configurations of resting-state networks in ADHD

**DOI:** 10.1101/734111

**Authors:** Neda Kaboodvand, Behzad Iravani, Peter Fransson

## Abstract

Attention deficit hyperactivity disorder (ADHD) is characterized by high distractibility and impaired executive functions. Notably, there is mounting evidence suggesting that ADHD could be regarded as a default mode network (DMN) disorder. In particular, failure in regulating the dynamics of activity and interactions of the DMN and cognitive control networks have been hypothesized as the main source of task interference causing attentional problems. On the other hand, previous studies indicated pronounced fluctuations in the strength of functional connections over time, particularly for the inter-network connections between the DMN and fronto-parietal control networks. Hence, characterization of connectivity disturbances in ADHD disorder requires a thorough assessment of time-varying FCs. In this study, we propose a dynamical systems perspective to assess how the DMN over time recruits different configurations of network segregation and integration. Specifically, we were interested in configurations for which both intra- and inter-network connections are retained, as opposed to commonly used methods which assess network segregation as a single measure. From resting-state fMRI data, we extracted three different stable configurations of FC patterns for the DMN, namely synergies. We provide evidence supporting our hypothesis that the ADHD brain differs in both recruitment rate and the topology of specific synergies between resting-state networks.

## 1. Introduction

Temporal fluctuations in functional interactions in the brain are structured to achieve an efficient balance between segregation and integration across various neural dynamical processes (Betzel et al., 2016; Zalesky et al., 2014). The dynamic nature in cooperative patterns in the brain brings up the possibility of controlling access to cognitive resources (Gu et al., 2015), which is in turn based on achieving a stabilized tradeoff between minimized metabolic costs and a highly optimized information processing strategy (Zalesky et al., 2014). This intention-specific stability can be facilitated through implementing different configurations of stable cooperative patterns between neural systems, phenomena which can be referred to as synergies. Generally speaking, a synergy can be described as a group of elements working together towards a common aim while assuring the stability of an outcome variable (Falaki et al., 2014; Kaboodvand et al., 2013; Latash et al., 2007; Schoner, 1995).

Given this background, it seems plausible that a neuroimaging analysis approach that is capable of providing a fine-grained temporal and spatial account of the dynamic nature of interactions between networks stands a good chance to adequately characterize and quantify recruited synergies in the brain. The aim of the present paper was two-fold. First, we present a novel approach to characterize synergies in brain configuration dynamics that is based on dynamical systems theory. Second, we show that our method can be applied to characterize and differentiate properties of brain synergies in healthy controls and an ADHD cohort.

Our proposed approach is based on a recurrence quantification technique (Marwan et al., 2007), together with a parameterization of resting-state fMRI brain connectivity data with the aim to create a state-space model of brain connectivity topology. Here, since we were interested in characterizing brain synergies in ADHD, we focused our efforts on the core neurocognitive networks, namely the default mode network (DMN) and the cognitive control networks (fronto-parietal networks (FPNs) and cingulo-opercular network (CON)) (Menon, 2011), which are systematically engaged during attention-demanding tasks (Menon, 2011). In particular, mounting evidence attests that ADHD could be regarded as a DMN disorder (Broyd et al., 2009; Castellanos and Proal, 2012; Sonuga-Barke and Castellanos, 2007; Uddin et al., 2008), considering DMN abnormalities as the main source of task interference causing attentional problems (like fluctuating attention and occasional periodic lapses in attention). Unsuccessful suppression of the DMN in transition from rest to task (Sonuga-Barke and Castellanos, 2007), as well as a low level of intra-DMN FC (Castellanos et al., 2008; Uddin et al., 2008) and failure in regulation of signaling between the DMN and FPNs (Sripada et al., 2014) have been suggested as the characteristic atypical functional connectivity patterns observed in ADHD. Notably, converging evidence from a number of brain imaging studies suggests a critical role for the CON in regulating the activity and interactions of the DMN and FPNs (Di and Biswal, 2014; Menon and Uddin, 2010; Sevinc et al., 2017; Sridharan et al., 2008), highlighting the necessity of a comprehensive method for studying these intrinsically coupled functional networks.

Importantly, the highly variable nature of inter-network connections, particularly linking the DMN and FPNs over time (Zalesky et al., 2014), have not been adequately taken in to account in the previous ADHD studies. Moreover, it has been shown that patients diagnosed with ADHD have a higher variability in their resting-state BOLD signals compared to controls (Mowinckel et al., 2017; Nomi et al., 2018). Therefore, the main aim of the present study was to investigate group differences between healthy controls and AHDH for the occurrence rate and topology (or configuration) of stable and synergetic cooperation patterns for the DMN.

## 2. Materials and Methods

### 2.1. Data used and data pre-processing

We used the resting-state fMRI data from healthy controls (n = 121, age range: 21-50 yr) and subjects diagnosed with ADHD (n = 40, age range: 21-50 yr) from the University of California LA Consortium for Neuropsychiatric Phenomics study. Further details regarding the cohorts can be found in (Poldrack et al., 2016). The standard image pre-processing steps, including realignment and unwarping, slice-timing correction, segmentation, normalization into Montreal Neurological Institute (MNI) template space, and smoothing using a Gaussian kernel with a full-width at half-maximum (FWHM) of 6 mm, were carried out using the Statistical Parametric Mapping software (SPM12; Welcome Department of Cognitive Neurology, University College London, London, United Kingdom). Three healthy participants with high (FD > 0.6) frame-wise displacement were excluded from further analysis (Power et al., 2012).

### 2.2. Identification of functional networks and time-varying inter-network functional connectivity

We employed independent component analysis (ICA) as implemented in the GIFT toolbox (Allen et al., 2011; Calhoun et al., 2001) to identify resting state networks (RSNs), with a particular focus on the DMN and cognitive control systems (FPN and CON). By applying ICA as a multivariate data-driven approach, the need for an arbitrary decision on regions of interest is alleviated. Using spatial ICA, we decomposed the fMRI data into spatially independent and temporally coherent component maps and their corresponding time-courses, representing the dynamics of the BOLD signals within every individual component (Calhoun et al., 2001).

In short, BOLD signals of each voxel was divided by its average intensity (i.e., intensity normalized) to increase accuracy and reliability of the ICA decomposition. Intensity-normalized data for all participants were concatenated across time. Subsequent to temporal concatenation, the ICA model order of 30 was estimated by applying the Minimum Description Length (MDL) criterion (Rissanen, 2004). The dimensionality of the data was reduced by applying two steps of data reduction using principal component analysis (PCA1: 100 components; PCA2: 30 components), followed by employing the Infomax algorithm (Bell and Sejnowski, 1995) for detecting temporally coherent networks. The ICA decomposition was repeated 20 times with different initial points to ensure stability of the estimated components. Finally, subject-level spatial maps and time-courses were estimated using the GICA3 back-reconstruction method (Allen et al., 2011; Kaboodvand et al., 2018).

One-sample t-tests were performed on the spatial maps, followed by excluding visually inspected artefactual components (e.g., vascular, ventricle and motion artifacts). With regard to our aim of investigating the dynamics of brain connectivity, we identified six ICA components that closely resembled the RSNs that are believed to be of central importance to cognitive processing in general and of particular importance for ADHD. The six identified ICA maps are shown in Figure 1 (for t-values > 20) and they encompass the anterior and posterior subsystems of the DMN (aDMN and pDMN), and the posterolateral parietal subsystem of the DMN, centered at temporoparietal junction (TPJ), as well cognitive control related networks, including left and right FPNs (lFPN and rFPN) and the CON network (which may also be referred as the salience network).

**Figure 1.**
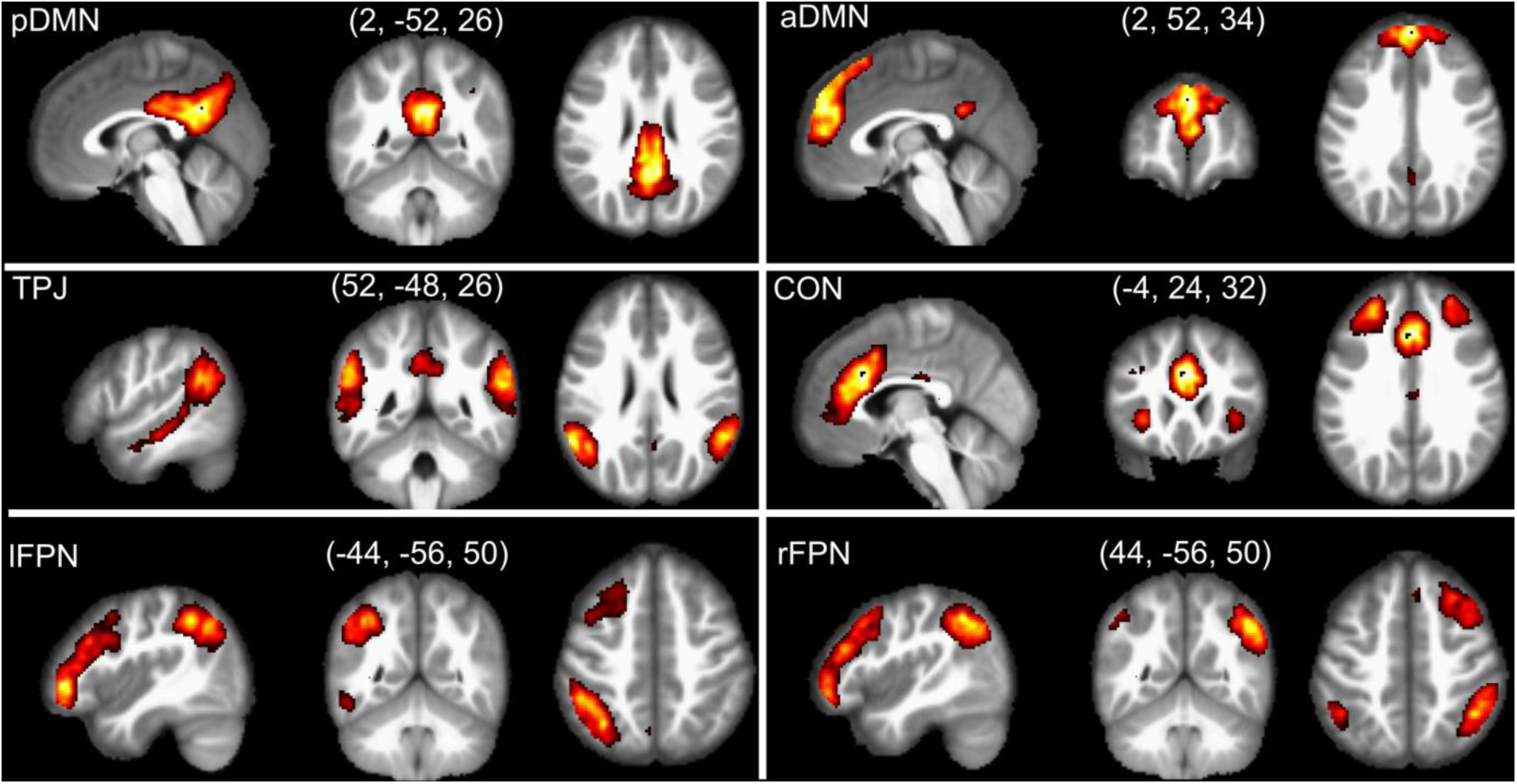
The six resting-state networks (independent-component analysis (ICA, t>20)) on concatenated fMRI data from all subjects) investigated in the present study include the posterior default mode network (pDMN), the anterior DMN (aDMN), posterolateral parietal DMN, centered at temporoparietal junction (TPJ), left and right frontoparietal control networks (lFPN and rFPN), and the cingulo-opercular network (CON).

Subject-level time-courses for all ICA components were detrended, despiked, and low-pass filtered (Butterworth, fifth-order, cut-off frequency equal to 0.15 Hz). There were no significant group differences in static functional connectivity (FC; z-transformed correlation coefficients) between any of the DMN subsystem and the cognitive control networks.

Next, time-varying FC was computed by measuring the instantaneous phase synchrony between ICA components (a detailed description is given in (Kaboodvand et al., 2019)). The instantaneous phase synchrony analysis is commonly used to estimate time-resolved FC/synchrony (Glerean et al., 2012; Ponce-Alvarez et al., 2015), because it provides single time-point resolution of time-resolved connectivity, with no need for an arbitrary choice of window length in contrast to correlation-based sliding window analysis. An illustration of the steps taken in the FC analysis is shown in Figure 2A.

**Figure 2.**
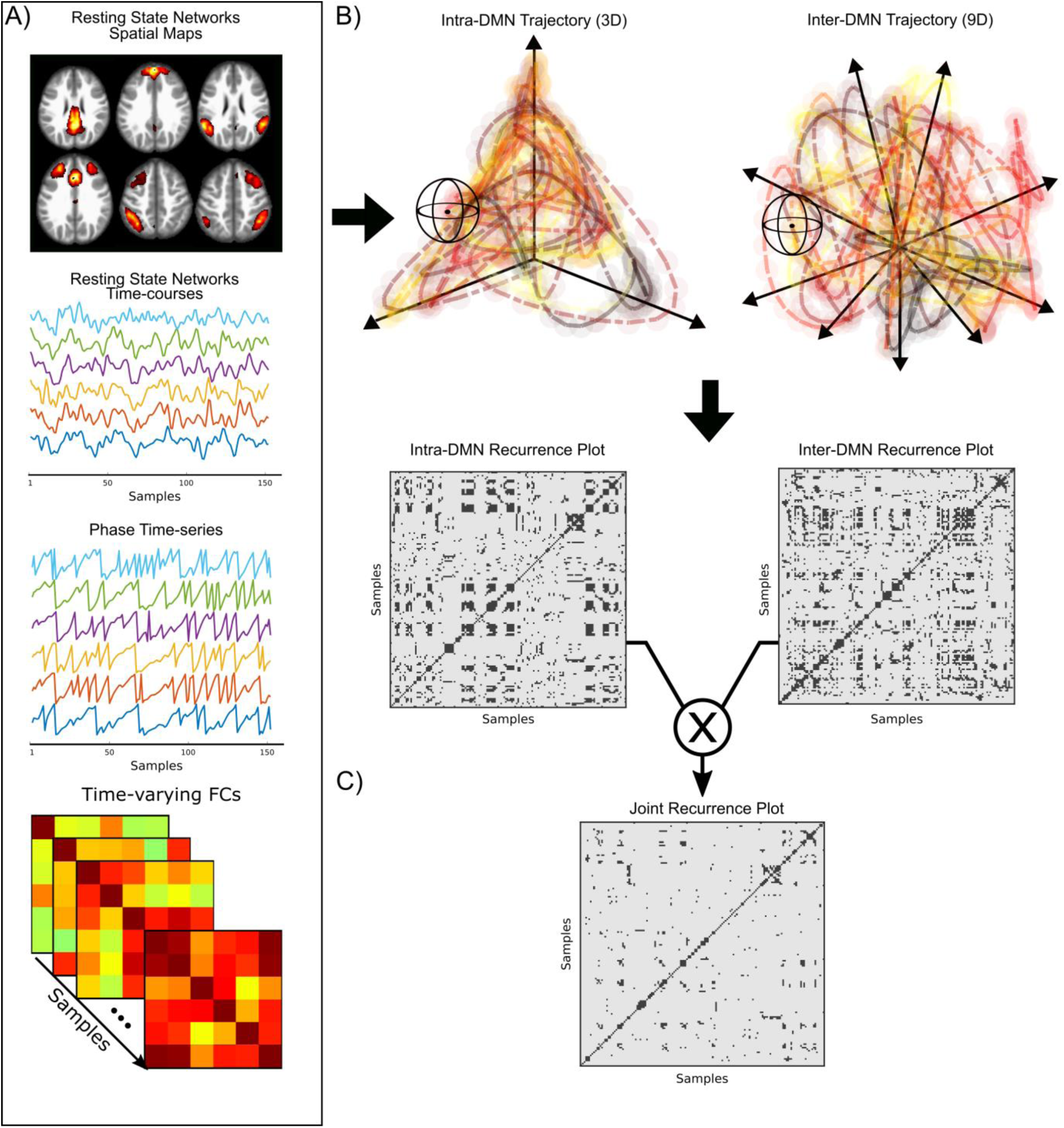
Illustration of the analysis pipeline. We used independent-component analysis (ICA) on resting-state functional MRI (fMRI) data to identify six resting-state networks and their respective time-courses. We applied the Hilbert transformation to the ICA time-courses to derive the analytic representation of each component’s signal, in order to calculate their respective instantaneous phase. The instantaneous functional connectivity (FC) between each pair of networks was defined by the cosine similarity of the phases obtained from the resting-state network signals (A). Next, we separately constructed the state-space trajectories for the intra-DMN (3-dimensions) and inter-DMN FCs (9 dimensions), followed by computing their respective recurrence plots. Note that the 3 dimensional projection of inter-DMN trajectory is plotted in hypothetical 9-dimensional space for illustration purposes (B). Finally, we computed the joint recurrence matrix from the recurrence plots built for the intra-DMN and inter-DMN FCs state-space trajectories (C).

### 2.3. Construction of state-space trajectories for network interactions

Dynamical systems theory is a mathematical framework that describes the behavior of the complex systems which evolve in time, and it relies on the concept of a state-space (also known as phase-space), that is the collection of all possible states of the system. Every possible state corresponds to one unique point in state-space, and the temporal evolution of the system in state-space traces a path which is called the “state-space trajectory”. Here, we apply a dynamical systems perspective to assess how individual brain networks interact with each other across time, all the while the configurations of both intra- and inter-network (in this study, the DMN) connectivity are retained, which is in contrast to previously used reductionist methods of computing network segregation as one single measure averaged across the whole scan (Chan et al., 2014), or as a single time-varying measure (Fransson et al., 2018; Fukushima et al., 2018).

Typically, when constructing state-space vectors, one is confronted with the problem of a time-varying measurement of only one observable system variable, since most often not all system variables are measurable. A frequently used approach to address this problem is the time-delay technique for state-space reconstruction (Takens, 1981), which is based on the time-delayed values of the observed system variable. However, in this study, we propose a method that treats network FC time-courses as independent system state-space variables.

Our topological state-space analysis of brain dynamics was based on the FC time-courses for the six resting-state networks as shown in Figure 1. As alluded to in the introduction, we were specifically interested in differences in synergies of networks that according to the previous literature are related to ADHD. We therefore focused our analysis on the intra-DMN (e.g. aDMN – pDMN) and inter-DMN (e.g. aDMN – lFPN) network interactions and their stable cooperation patterns (i.e. synergies). For each subject, intra- and inter-network state-space trajectories were constructed for the network whose segregation pattern is of interest (here: the segregation of DMN in respect to cognitive control networks).

Our formulation of fMRI brain connectivity into a topological state-space analysis can be described as follows. We can write the time-course of FC between two networks as *u*_*i*_ = *u*(*i*Δ*t*), where *i* = 1,…,*N* and Δ*t* is the sampling rate of the measurement. For the given system, the corresponding vectors in the state-space can be constructed from the time series of FCs by employing the generic functions:

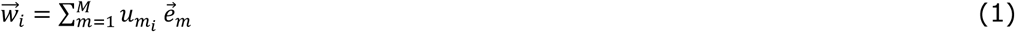

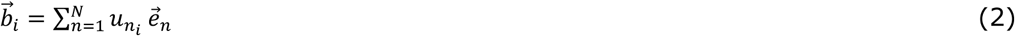

where 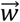 and 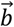 are intra- and inter-network state-space trajectories. M is the dimension of intra-network state-space (the number of intra-network connections). Similarly, N is the dimension of the inter-network state-space (the number of inter-network connections). In our case that pertains a focus on the DMN network and its relationship to the cognitive control networks, the dimensionality of intra-DMN state-space is N = 3 and the dimension of the inter-DMN state space is M = 9. The vectors 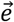 are unit vectors spanning an orthogonal coordinate system. Thus, for every individual, we calculated two state-space trajectories. The trajectory for intra-DMN network interactions resided in a 3-dimensional state space that is spanned the degree of connectivity between the three DMN subsystems (aDMN – pDMN, aDMN – TPJ and pDMN – TPJ). Accordingly, the trajectory for inter-DMN network interactions resided in a 9-dimenasional state-space, formed by 9 network-network FCs (aDMN – lFPN, aDMN – rFPN, aDMN – CON, pDMN – lFPN, pDMN – rFPN, pDMN – CON, TPJ – lFPN, TPJ – rFPN, TPJ - CON). Of note, network interactions outside the DMN were not included in the analysis (lFPN – rFPN, lFPN – CON, rFPN – CON).

### 2.4. Recurrences and recurrence plots

A fundamental property of deterministic dynamical systems is that after some time, the system states revisit previous states within an arbitrary short distance (Ott, 2002; Poincaré, 1890). This is the governing principle behind the idea of using recurrence plots to infer properties of dynamical systems from state-space representations (Eckmann et al., 1987). Briefly, a recurrence plot (or equivalently, a recurrence matrix), allows us to determine how often the state-space trajectory for a given system is in close vicinity of states it has visited previously. Hence, we can compute the recurrence matrix to identify the time-points when the state-space trajectory 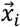 recurs in approximately the same area in state-space (Marwan et al., 2007). Recurrence points refer to the states which are in an ε-neighborhood of each other for some chosen value of the distance parameter ε. The recurrence matrix *R*_*i,j*_, is a binary matrix defined as:

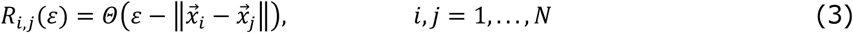

The Heaviside function is denoted as Θ(·) and the distance between two states (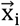 and 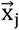) is computed by the distance (norm) function 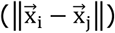. Here, we used the Euclidean distance (L2 distance) that has previously been commonly used for recurrence matrix quantification (Marwan et al., 2007). Importantly, a graphical representation of recurrence matrix, also known as a recurrence plot, provides the possibility to investigate properties of recurrences in a_n_ M-dimensional state-space trajectory through a two-dimensional graph. Recurrent and non-recurrent pairs of state vectors are illustrated as black (R_i,j_ = 1) and white (R_i,j_ = 0) dots in the recurrence plot respectively. In other words, a black dot at coordinate (i, j) represents a recurrence of the system’s state as denoted by 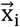 at time j (i.e.,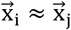). The graphical concept of recurrence plots in the context of brain network connectivity is illustrated in Figure 2B.

Notably, the values of the recurrence matrix are binarized based on an arbitrary thresholdε. So, we need to find a value of ε that is suitable for our purposes and it will to some extent inevitably be a compromise between different aims. If an excessively small threshold is selected, there may be almost no recurrence point recorded in the recurrence plots, whereas a too large threshold will introduce false positives. Indeed, the existence of noise in the measurements requires a reasonably large threshold in order to preserve relevant structures of recurrences in the data. In the literature, both absolute (fixed) and relative (fixed recurrence density) strategies have been suggested (Marwan et al., 2007), a problem that is akin to the ongoing discussions of how to select appropriate thresholds for graph theoretical analysis in neuroimaging (Fornito *et al.*, 2016). Common choices of the threshold parameter ε is to let it depend on a recurrence rate (equivalent to edge density) of 5% (Donner et al., 2010; Marwan et al., 2009; Schinkel et al., 2008; Zou et al., 2018), or 1% (Marwan et al., 2007). Since we were interested in comparing recurrence plots for healthy versus ADHD, we preferred to use an absolute threshold, because we expected a significant group difference in recurrence density, which when translated into a threshold value would result in significant inter-group difference in the values of the threshold ε. Therefore, after we separately constructed the state-space trajectories for the intra-DMN and inter-DMN FCs, we computed their respective weighted recurrence plots separately. Subsequently, we estimated the absolute threshold corresponding to the 1% density for the joint recurrence plots at subject-level, followed by averaging them across all subjects to find a common absolute threshold for all individuals.

### 2.5. Joint recurrence plots

We have at this point derived a method to compute state-space trajectories for both intra-and inter-DMN network-network interactions, but we additionally need a method to combine the information from both state-spaces into a single framework. A possible way to do this is to compute the joint recurrence plots, a multivariate extension of recurrence plots, which allows for studying the joint recurrences of different systems by examining the simultaneous occurrence of recurrences, while preserving their respective individual state-spaces (Marwan et al., 2007; Zou et al., 2018). The joint recurrence matrix is defined as the element-wise product of the individual recurrence matrices *R*^*p*^:

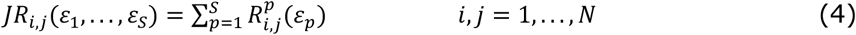

where the joint recurrence plot of *p* = 1,…,*S* systems, each of them thresholded at the *ε*_*p*_-level, is indicated by the *JR* matrix. Here, we computed the joint recurrence matrix from the recurrence plots computed separately for the 3-dimensional intra-DMN and the 9-dimension inter-DMN state-space trajectories. In our case, a recurrence will take place (JR_i,j_ = 1) if a point/state at *i* time in the intra-DMN trajectory returns to the neighborhood of a former point/state across that trajectory at time *j*, and at the same time a state at time *i* on the inter-DMN trajectory returns to the vicinity of a former state across inter-DMN trajectory at time *j* (Marwan et al., 2007; Zou et al., 2018). This implies that, focusing on laminar states (see definition below), we study the joint probability that a stable (for at least 2×TR = 4 seconds) configuration of intra-DMN FCs as well as a stable configuration of inter-DMN FCs, happen simultaneously. A schematic representation of the methodology described here is depicted in Figures 2B and 2C.

### 2.6. Quantifying recurrence in state-space

Recurrence matrices enable us to easily visualize and gain insights into the temporal evolution of state-space trajectories, specifically in a case like ours which contains a high number of dimensions. Importantly, typical patterns of recurrences are associated with particular types of system behavior (Marwan *et al.*, 2007). Recurrence points may form structures, classified into either diagonal lines, or vertical/horizontal lines, which are characteristics for different dynamical behaviors of the system under investigation (for examples of this behavior, see recurrence plots shown in Figure 2B). On the other hand, if trajectories in state-spaces fluctuate strongly, they may be revisited only for short periods of time which results in single points in the recurrence plot. Relatedly, the recurrence rate is the simplest measure derived from the recurrence plot that is simply the density of recurrence points. A vertical (or horizontal) line structure in the recurrence plot signifies a state that is trapped for some time. This is a property that is of central importance in the current study as states which do not change or change very slowly over time are an indication of laminar states (Marwan et al., 2007). Hence, laminar states in terms of the presence of vertical or horizontal lines in the recurrence plots suggest that the trajectories of network-network functional connectivity are in a stable configuration (i.e., compare with laminar flow).

Further, the length of a vertical (horizontal) line in time units (denoted by V) represents the duration in which the state does not change or changes very slowly (R_i,j +v_ = 1, ∀ v = {1,…, V}). The laminarity of the system is defined as the ratio of recurrence points that form vertical lines with the minimum length of V_min_ (commonly set to 2) in relation to all observed recurrence points. The laminarity alone does not provide us with any information about duration of stable states. However, we can get a handle on this by computing the average time interval that the system is trapped at particular states. This property is known as trapping time (Marwan *et al.*, 2007). Figure 3 provides an illustrative example of a laminar state observed for the intra-DMN and inter-DMN state-space trajectory in a randomly selected individual.

**Figure 3.**
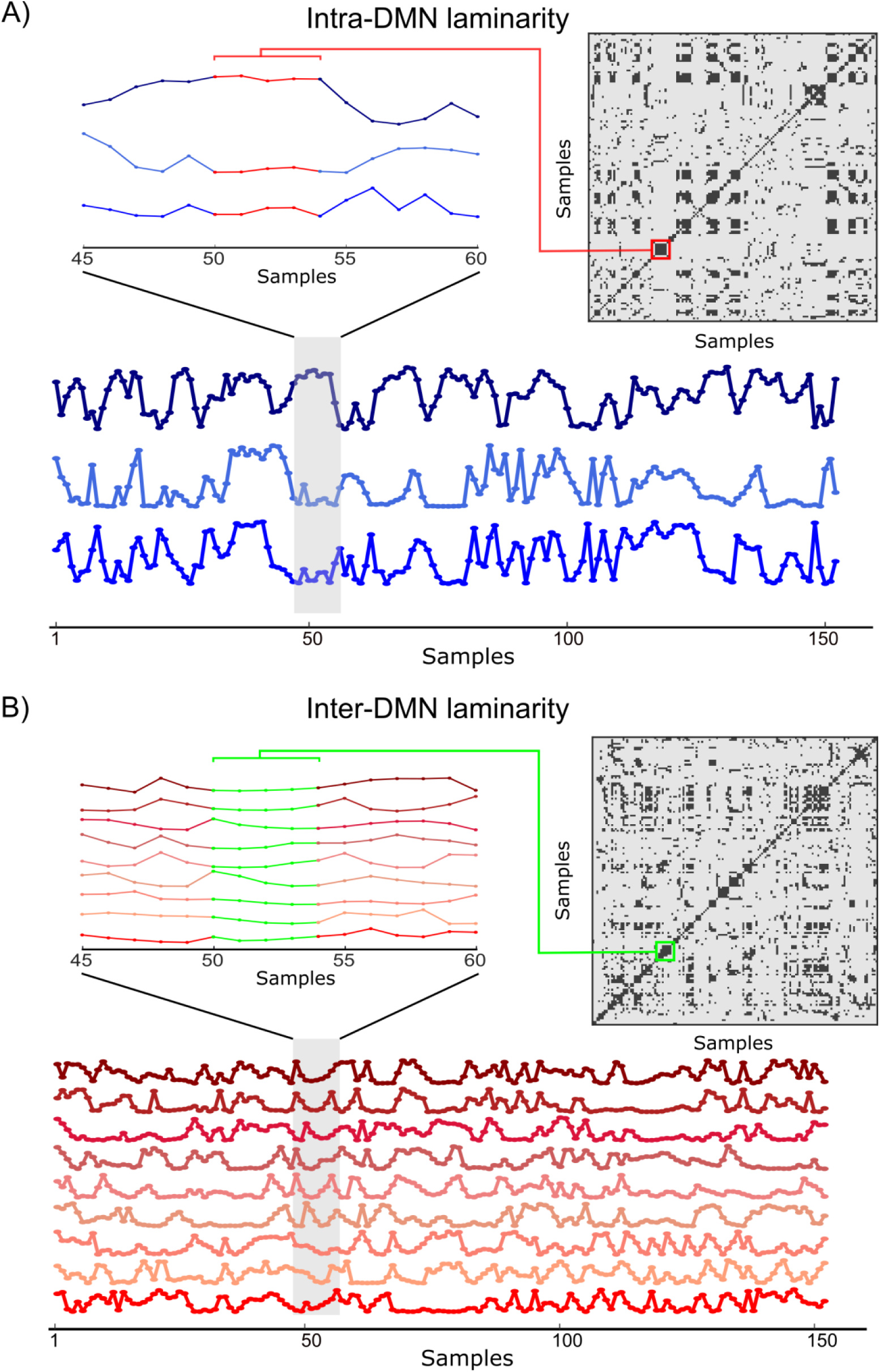
Illustrative example of laminarity in one arbitrarily chosen subject. The 3 underlying time-courses of intra-DMN trajectories as well as the 9 underlying time-courses of inter-DMN trajectory are illustrated in panel (a) and (b), respectively. We note that different patterns of connectivity may be present in each laminar state. For example, the laminar state of this example is associated with one strong and two weak intra-DMN connections.

We computed both laminarity and trapping time for the joint recurrence plot of the DMN (i.e. by combining data from the intra- and inter-DMN recurrence plots as detailed above). This was done at the subject-level, followed by regressing out age, sex and FD. We compared the average difference between the two groups (healthy and ADHD), by bootstrap resampling (10,000 samples). Subsequently, the 95% confidence interval was derived for detecting significant group differences in laminarity and trapping time.

### 2.7. Identification of network synergies

As a last step in our analysis, we aimed to simultaneously assess the behavior of time-varying intra-DMN FCs as well as the time-varying FCs of the DMN with cognitive control networks (aka inter-DMN FCs). This was done in order to achieve a complete picture of putative differences in the synergistic cooperation patterns of the DMN in ADHD compared to controls. In this context, we use the term synergy to refer to all instances in time where cooperative patterns of connectivity involving the DMN (as observed from both intra-DMN as well as inter-DMN trajectories) are simultaneously stable for a time-segment (i.e., laminar states in the joint recurrence plot) that lasts least 4 seconds (2×TR). Accordingly, we wanted to test the hypothesis that ADHD can be traced to quantitative differences in parameters that details synergies between brain networks, such as occurrence rate and trapping time.

As a starting point to test our hypothesis, we compared recurrence structures in the joint recurrence plots (with the particular focus on vertical/horizontal structures), constructed from state-space trajectories of the intra-DMN and inter-DMN FCs. Next, we assessed the degree of cooperation patterns between networks during stable/laminar states. To do this, we averaged the state-space trajectories for time-points associated with the occurrence of all laminar states, as they have almost constant values during a laminar state. Hence, every laminar state was correspondingly characterized by a vector consisting of 12 FC measures (3 intra-DMN FCs + 9 inter-DMN FCs). Next, we concatenated all the 12-dimensional connectivity vectors associated with the respective laminar states, across all subjects, in order to split connectivity vectors into k clusters, representing different synergies recruited by the DMN. In general, clustering of data is usually associated with several choices that need to be made, such as choosing the number of clusters and selecting the single best partition among different runs that the clustering algorithm is run. We here opted to use consensus clustering (Ghosh and Acharya, 2011; Vega-Pons and Ruiz-Shulcloper, 2011) since it attempts to alleviate some of aforementioned problems by detecting common structures of an ensemble of partitions, although it requires an arbitrary threshold for the consensus rate. However, hierarchical consensus clustering (Jeub et al., 2018) allows us to get the most stable partitions, without requiring us to decide on the exact number of clusters or even the threshold for consensus clustering.

In detail, we first applied the Calinski-Harabasz clustering evaluation criterion (Calinski and Harabasz, 1974) to the 12-dimensional FC vectors calculated for all subjects, suggesting that an optimal number of clusters was 2. Second, we used the k-means clustering method (1000 repetitions), to create an ensemble of partitions for the consensus clustering step. Third, we employed a hierarchical consensus clustering procedure to identify a consensus clustering at a significance level alpha = 0.01 (Jeub et al., 2018). By using the hierarchical consensus clustering approach, we obtained 3 clusters, for which the mean vector (centroid) for each cluster served as a prototype for each synergy.

We compared the control versus the ADHD group with regard to the recruitment rate of every detected synergy. We here define the recruitment rate of an individual synergy by computing the sum of time-intervals (consisting of consecutive time-points), which their respective connectivity patterns were categorized as members of that particular synergy.

Additionally, we investigated if the ADHD cohort not only differed in terms of their recruitment rate, but also differed in the synergetic cooperation patterns (i.e., the topology of synergies). Hence, we compared the average difference of the 12-dimensional connectivity vectors between the two cohorts, by bootstrap resampling (10000 samples). A 95% confidence interval was used for detecting significant group differences in synergistic cooperation patterns among brain networks.

## 3. Results

Based on the data obtained the joint recurrence plot (JR) which was calculated from recurrence plots that combined intra-DMN and inter-DMN state-space trajectories, we observed a significantly higher degree of mean laminarity (CI: [0.67 5.23]) and mean trapping time (CI: [0.02 0.09]) for the healthy compared to the ADHD cohort (Figures 4A and B).

**Figure 4.**
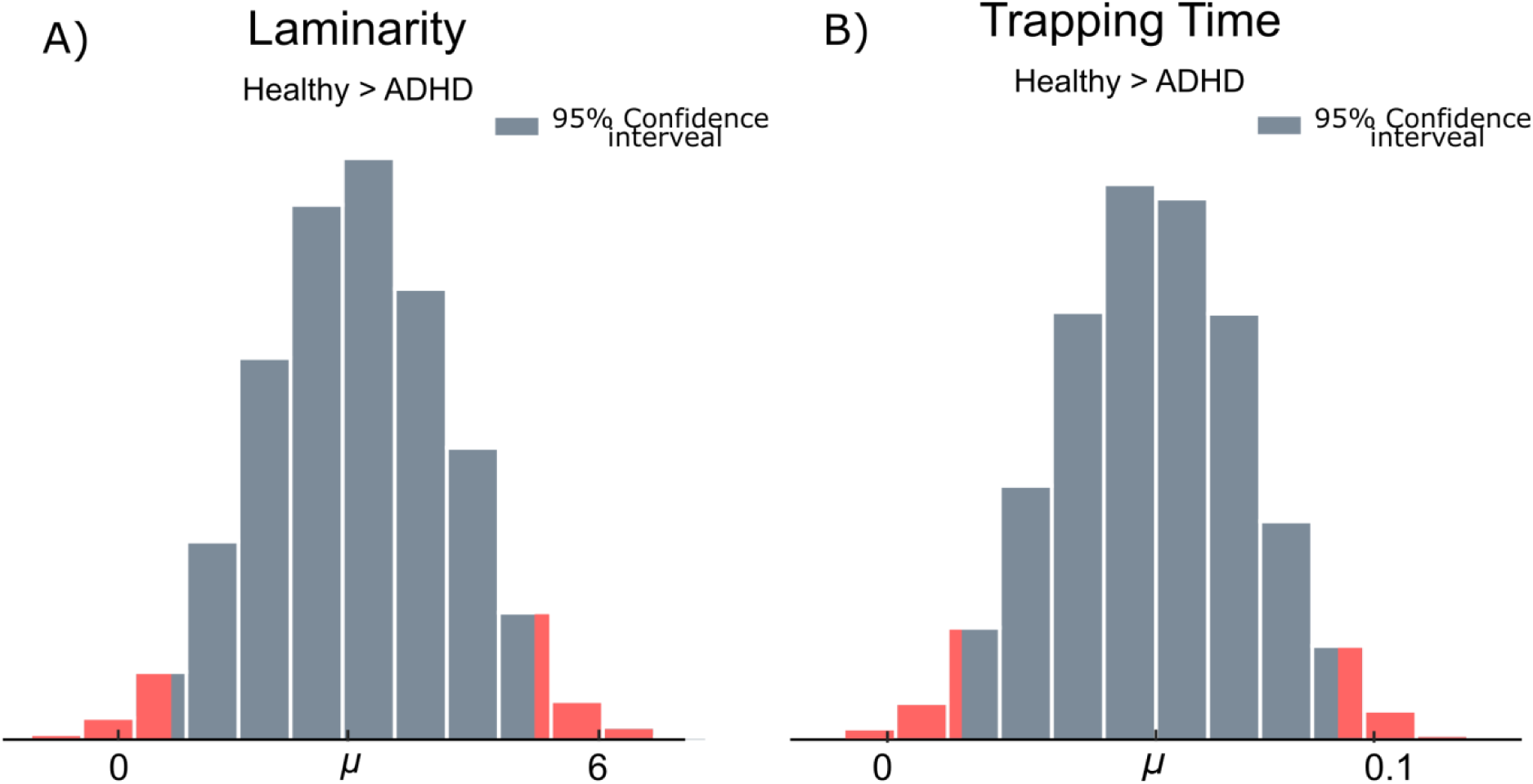
Statistical resampling tests showed a higher degree of mean laminarity (A) as well as mean trapping time (B) in the healthy compared to the ADHD cohort. Age, sex and FD were regressed out. The average group-difference in each measure was calculated by bootstrap resampling (10000 samples). The 95% confidence intervals are colored in gray.

The results shown in Figure 4 suggest that when all intra-as well as inter DMN network-network interactions are considered collectively, a lesser degree of stable cooperation between networks was found in the ADHD cohort. In other words, when averaged across all possible synergies (3 synergies), we find that periods of stable cooperations (as captured by the degree of laminarity) occur significantly less frequently in ADHD than controls (Figure 4A). Moreover, we show that the corresponding periods of stable cooperations are shorter in duration (i.e. shorter trapping time) in ADHD than controls (Figure 4B).

Next, we sought to quantify the relative contribution of stable cooperation from each individual network-network pair. Accordingly, our investigation of synergistic cooperation among DMN and cognitive control networks was focused on the time-intervals for which the configurations of both intra- and inter-DMN connections have jointly reached their steady states. To do this, we applied a consensus hierarchical clustering approach to the 12-dimensional connectivity vectors derived for all laminar states, concatenated across all subjects, which resulted in three clusters (synergies). Of note, each synergy provides detailed information regarding the relative values of the 12 DMN-related FCs, when both intra-DMN and inter-DMN trajectories are simultaneously invariable for a time interval with a duration of at least 4 seconds. The results are shown for both groups in form of radar charts in Figure 5.

**Figure 5.**
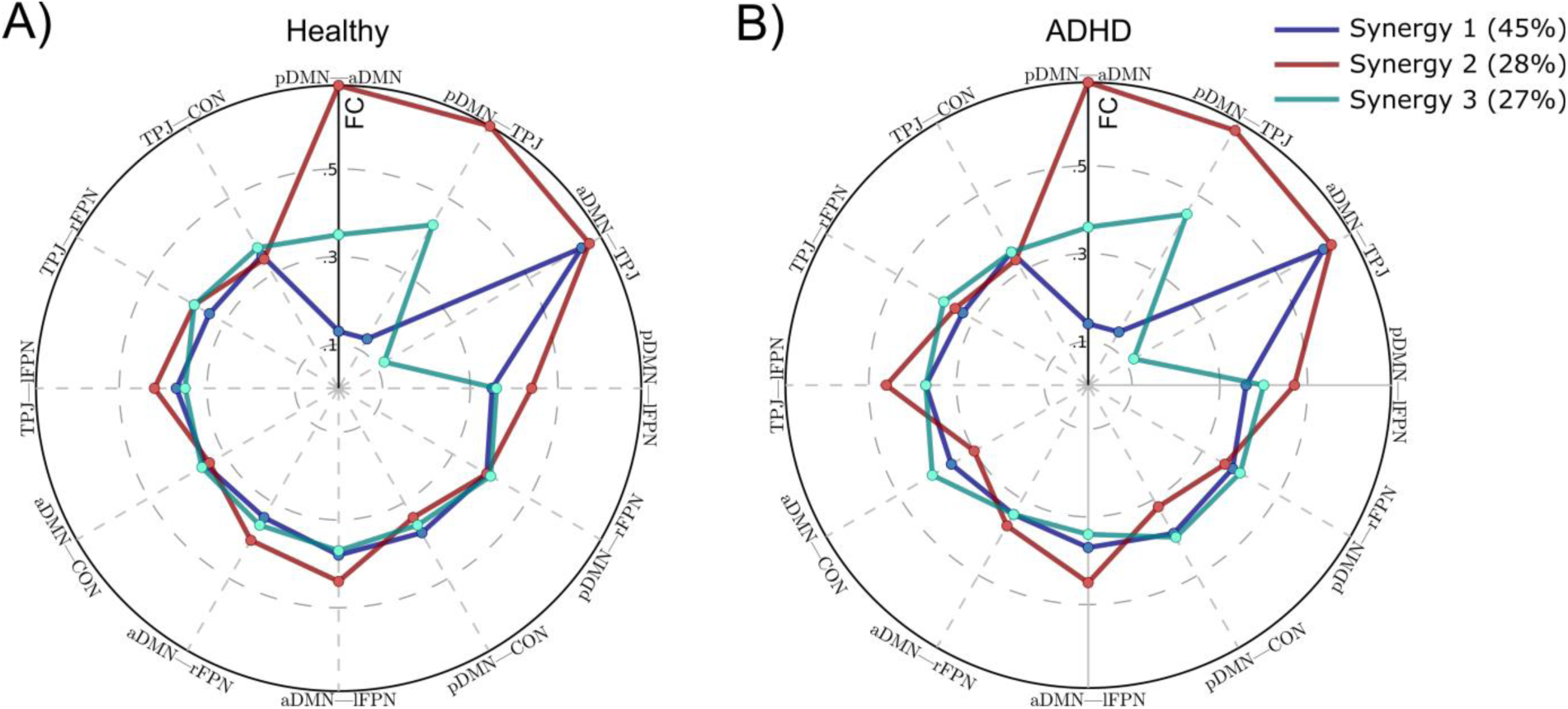
Group-level assessment of the average degree of brain network synergies recruited during laminar states for state-space trajectories that span both intra-DMN and inter-DMN network-network interactions. For each synergy the percentage of all laminar states which were classified as belonging to that particular synergy are shown in the parenthesis. For abbreviations and anatomical description of the six networks, see Figure 1.

A visual inspection of the results shown in Figure 5 suggest an overall agreement of synergies in both groups, however some small differences between groups are also apparent. In detail, the first synergy (labeled as #1 in Figure 5) is characterized by a marked decrease in FC between the pDMN and the other two DMN sub-networks (aDMN and TPJ), but also a quite strong degree of FC between the aDMN and TPJ. The other nine network pairs show a moderate level of connectivity. We note that 45 percent of all laminar states were classified as belonging to synergy #1, which means that it is the most commonly recruited configuration of cooperation between networks in our model. Interestingly, the second synergy (#2) is foremost defined by strong levels of cooperations between all subsystems of the DMN (aDMN, pDMN, TPJ). In contrast, synergy #3 display evenly distributed cooperations among all network pairs with the exception of the aDMN-TPJ cooperation that was very small. The second and third synergies were recruited for 28 and 27 percent of all laminar states, respectively.

It is important to point out that all three synergies depicted in Figure 5 were recruited by all subjects in both groups, but as stated previously, on average, healthy subjects recruit these synergies more frequently across time (higher degree of laminarity) (Figure 4A). Moreover, when healthy subjects have on-going synergistic cooperation between networks, they keep them for a longer period of time (longer trapping time) (Figure 4B).

Next, we were interested in the question of whether the presence and duration of the three synergies shown in Figure 5 were statistically different between cohorts. When comparing the healthy cohort with the ADHD cohort, we found a significantly stronger recruitment rate (computed as the sum of all time-intervals with stable (laminar) cooperation between networks) for the second synergy (CI: [0.23 7.04]). The distribution of differences between the healthy and ADHD cohorts in terms of recruitment rate for the second synergy is shown in Figure 6.

**Figure 6.**
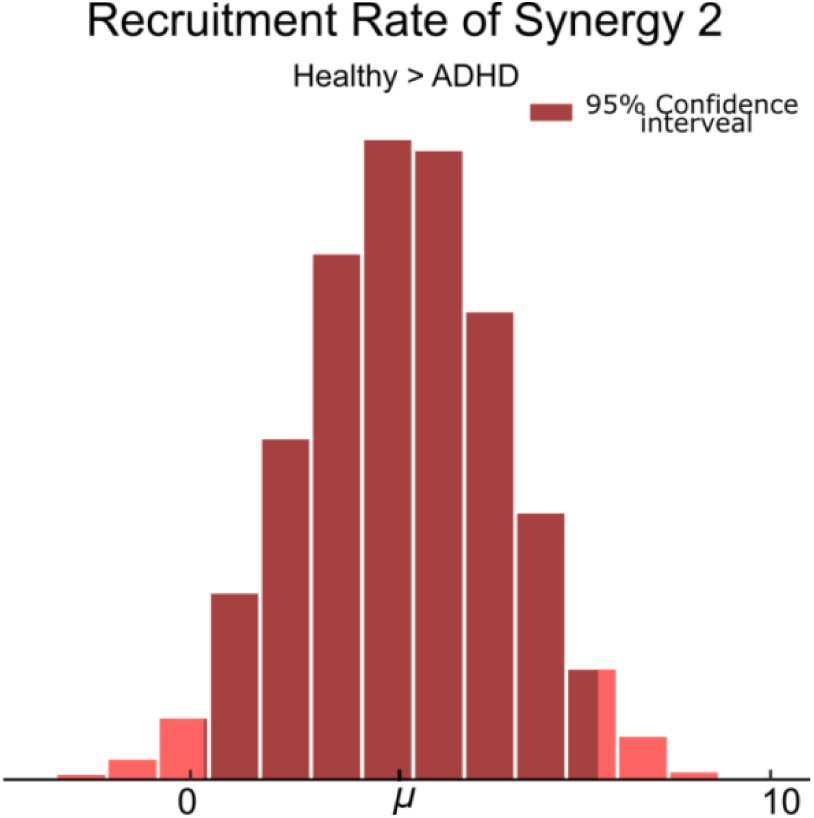
Statistical resampling test (10000 samples) showed a stronger recruitment rate for synergy 2 (computed as the sum of the durations for all time-intervals with stable (laminar) cooperation between networks) in the healthy cohort compared to the ADHD cohort (CI: [0.10 6.82]). Of note, the second group-level synergy (#2) is associated with strong within-DMN connectivity (see also Figure 5).

Differences between the ADHD and healthy cohorts in terms of synergetic cooperation patterns (i.e., the topology of synergies) were foremost found in synergy #3. Healthy controls showed a significantly weaker inter-DMN connectivity than ADHD group, including the pDMN — lFPN (CI: [-0.076 -0.005]), pDMN — CON (CI: [-0.074 -0.002]) and aDMN — CON (CI: [-0.089 -0.017]) network pairs.

Additionally, the second synergy (which had less recruitment rate for the ADHD group) showed a marginally lower level of coherence between the DMN sub-networks and the lFPN (pDMN — lFPN, CI: [-0.074 0.004]; aDMN — lFPN, CI: [-0.055 0.022]; TPJ — lFPN, CI: [-0.079 0]) in the healthy cohort compared to ADHD, but a trend for stronger level of coherence for the DMN sub-networks with rFPN and CON (pDMN — rFPN, CI: [-0.007 0.072]; aDMN — rFPN, CI: [-0.002 0.079]; TPJ — rFPN, CI: [-0.004 0.77]; pDMN — CON, CI: [-0.017 0.061]; aDMN — CON, CI: [-0.007 0.069]; TPJ — CON, CI: [-0.026 0.052]).

## 4. Discussion

We postulate that different brain disorders may manifest themselves by either disturbed topology of synergies, or their respective occurrence rates (i.e., how often a particular synergy is being recruited).

Using a topological state-space representation of an individual network’s connectivity patterns, we have shown that different constellations of brain connectivity across time can be described in a framework of synergies in the form of temporary but stable cooperations between multiple networks in the brain. In the present study of putative abnormalities in connectivity in ADHD, we focused on intra-as well as inter-DMN functionality. In this context, we used the term synergy (steady-state synergy) to refer to any particular stable cooperation pattern taken by the DMN, which is recruited when both intra-DMN and inter-DMN trajectories are simultaneously invariable for a time interval with at least 4 seconds duration (i.e., laminar state of the joint recurrence plot, lasting at least 4 seconds (2×TR)).

We observed an overall reduction in synergetic cooperation between networks, both in terms of frequency and duration of laminarity states (stability of patterns of cooperativity) in ADHD compared to healthy controls (Figures 4). This finding is in line with the previous literature that have found a higher variability of the resting-state BOLD signals for the individuals with ADHD (Mowinckel et al., 2017; Nomi et al., 2018).

Interestingly, we found that the steady-state configuration associated with the segregation of the pDMN network from the aDMN and TPJ networks together with a strong connection between the aDMN and TPJ networks, was the most predominant synergy during resting-state since almost half of all stable states were assigned to synergy #1 (Figure 5). Speculatively, synergy #1 might reflect an inner organization of the DMN that is often recruited and thus beneficial for performing specialized functions within the DMN. Noteworthy, we found no significant group differences in the synergistic cooperation patterns (i.e., the topology) of synergy #1, except for a positive trend of a stronger aDMN—TPJ FC in the healthy group (CI: [0.001 0.05]). It also deserved to be mentioned that there was a trend of longer trapping times for synergy #1 in the healthy cohort compared to ADHD (CI: [0 0.17]). Relatedly, the aDMN and TPJ networks have previously been suggested as networks involved in social cognition, and they are frequently co-activated in studies about theory of mind or mentalizing (Frith and Frith, 2006), such as affective speech comprehension (Frith and Frith, 2006; Hervé et al., 2012; Saxe, 2006). Indeed, functional integration between the aDMN and TPJ network seems to be beneficial for social cognition, and the smaller average dwell time in the laminar states recruiting this synergy as well as the smaller aDMN - TPJ cooperation for synergy #1 may be indicative of impaired social cognition that is often observed in ADHD (Uekermann *et al.*, 2010).

The overall topology of synergy #2 was across groups marked by an overall strong cooperation within the subsystems of the DMN. This finding suggests that the brain needs a considerable amount of time intervals during which it can exchange information within the DMN. Interestingly, we found a significant lower recruitment rate of synergy #2 in ADHD compared to controls (Figure 6), which suggests that the degree of cooperative patterns of connectivity within the DMN could be interpreted as a biomarker for ADHD dynamic connectivity. Although the differences between groups for individual network pairs were not significant (i.e., no significant topology difference), we could observe a trend for a lower degree of coherence between all three subsystems of the DMN and lFPN in the healthy cohort compared to ADHD. On the other hand, we note a trend for a stronger degree of coherence between all three DMN subsystems and both rFPN and CON.

It should be pointed out that time-resolved fMRI connectivity by instantaneous phase synchrony analysis as employed in this study, is only comparable to the absolute values of correlation-based sliding window analysis in the sense that strong positive/negative correlations display high phase synchrony (Pedersen et al., 2018). Therefore, stronger coherence between the DMN and rFPN as observed here, may be translated into either stronger positive correlation between these networks, or more anti-correlation between them (see also (Fransson, 2005)).

It has been previously suggested that the frontoparietal regions involved in stimulus-driven orienting and directing attention to salient events, are largely lateralized to the right hemisphere (Allen *et al.*, 2011; Corbetta and Shulman, 2002; Vincent *et al.*, 2008), whereas the left frontoparietal regions are particularly important for explicit memory retrieval (Iidaka *et al.*, 2006). On the other hand, the CON (also referred as salience network) is the other cognitive control network, anchored in the anterior insula and dorsal anterior cingulate cortex (see also Figure 1), which is primarily known as being involved in integrating sensory data with autonomic, visceral and somatic markers, in order to identify the biologically and cognitively relevant stimuli (salient events), particularly to facilitate attentional reorienting and access to memory resources in high-level cognitive control and attentional processes (Menon, 2011; Seeley et al., 2007; Sevinc et al., 2017; Sridharan et al., 2008). Additionally, these cingulo-opercular regions’ activity regulates the activity and interactions of the other large-scale networks (Di and Biswal, 2014; Menon and Uddin, 2010; Sevinc et al., 2017). Therefore, the interaction between the rFPN and CON has a key role in the hierarchical initiation of cognitive control signals (Di and Biswal, 2014; Menon and Uddin, 2010; Sevinc et al., 2017). Hence, we further looked into the coherence between rFPN and CON, during the intervals associated with strong within-DMN integrity (i.e., second synergy). We found that Synergy #2 in the ADHD group has significantly stronger coherence between these two key control networks (rFPN — CON, CI: [-0.092 -0.017]). A similar pattern was observed for Synergy #3 (rFPN — CON, CI: [-0.07 0.004]), although it was not significant. We speculate that higher ADHD-related coherence between rFPN and CON, during the intervals associated with strong within-DMN integrity, may lead to the interference of irrelevant information causing distraction. Importantly, there was an opposite trend (rFPN — CON, CI: [-0.003 0.053]) for the first synergy, associated with low DMN integrity.

A prominent feature of Synergy #3 was a weak degree of cooperation between the aDMN and the TPJ subsystem of the DMN. When we further looked into the group-effect on the topology of this synergy, we observed that the activity of pDMN was less coherent with both lFPN and CON. Additionally, while the aDMN was less coherent with CON in the healthy group, there was a trend for more coherent activity with the lFPN. There were exactly opposite trends for the TPJ, so that it showed a tendency for higher coherence with CON, but lower coherence with the lFPN.

In the current study we opted to focus on the time-segments during which a stable (lasting at least for 4 seconds) configuration of intra-DMN FCs as well as a stable configuration of inter-DMN FCs, happen simultaneously, and subsequently the overall configuration of stable cooperation patterns recruited by the DMN at those intervals was termed synergy (or steady-state configuration). Specifically, we tested our hypothesis that abnormalities in ADHD brain connectivity are linked to the occurrence rate and the topology of DMN synergies, which are steady-state configurations. However, the important role of transient configurations of connections for the balance between information exchange and metabolic costs has previously been shown (Zalesky et al., 2014). Therefore, future studies with higher temporal resolution may focus on the transient configurations, or even a combination of both transient and steady-state configurations, although this may be of less interest for ADHD studies.

We propose a dynamical systems perspective to assess how a brain network (in this study, the DMN) takes different configurations of segregation and integration in time, notably by preserving the configurations of both intra- and inter-network connections, as opposed to commonly used methods which assess the network segregation computed as one single measure. We extracted different stable configuration patterns for the network under study, namely synergies. We provided some evidence supporting our hypothesis that the ADHD brain differs in both recruitment rate and the topology of specific synergies.

## Acknowledgement

P.F. was supported by the Swedish Research Council (grant No. 2016-03352) and the Swedish e-Science Research Center.

